# Modulation of input to the spinal cord; contribution of GABA released by interneurons and glial cells by polarization at the entry of sensory information

**DOI:** 10.1101/2024.04.04.588053

**Authors:** Ingela Hammar, Elzbieta Jankowska

**Affiliations:** Dept. of Neuroscience and Physiology, Sahlgrenska Academy, University of Gothenburg, Sweden

**Keywords:** Myelinated nerve fibres, excitability, refractory period, GABA, glial cells, rat

## Abstract

Modulation of input from primary afferent fibres has long been examined at the level of the first relay neurons of these fibres. However, recent studies reveal that input to the spinal cord may also be modulated before action potentials in intraspinal collaterals of afferent fibres reach their target neurons, even at the level of the very entry of afferent fibres to the spinal grey matter. Such modulation greatly depends on the actions of GABA via extrasynaptic membrane receptors. In the reported study we hypothesized that the increase in excitability of afferent fibres following epidural polarization close to the site where collaterals of afferent fibres leave the dorsal columns is due to the release of GABA from two sources: not only terminals of GABAergic interneurons but also glial cells. We present evidence, primo, that GABA from both these sources contributes to a long-lasting increase in the excitability and a shortening of the refractory period of epidurally stimulated afferents fibres and, secondo, that effects of epidural polarization on GABA-containing terminals of GABAergic interneurons and on glial cells are more critical for these changes than direct effects on the stimulated fibres. The experiments were carried out in deeply anaesthetized rats in which changes in compound action potentials evoked in hindlimb peripheral nerves by dorsal column stimulation were used as a measure of the excitability of afferent fibres. The study throws new light on the modulation of input to spinal networks but also on mechanisms underlying the restoration of spinal functions.

## Introduction

Modulation of input from primary afferent fibres has long been examined at the level of the first relay neurons of these fibres by presynaptic inhibition or by changes in the excitability of their target neurons However, recent studies reveal that input to the spinal cord may also be modulated at the level of entry of afferent fibres to the spinal grey matter (see Lucas-Osma *et al*., 2018; Hari *et al*., 2022). Such modulation involves a long-lasting increase in fibre excitability for which there are strong indications that it is GABA-dependent (Li *et al*., 2020). In the reported study we hypothesized that the effects of GABA on the branching regions of the afferent fibres at the site where their collaterals exit the dorsal columns are secondary to the direct effects of epidural polarization on afferent fibres as well as to modulation of the release of GABA around the fibres. GABAergic interneurons and glial cells were both considered as sources of polarization-dependent release of GABA and we present evidence that both may contribute to the long-lasting effects of epidural polarization.

The polarization evoked increase in the excitability combined with the shortening of the latency of activation of afferent fibres (Li *et al*., 2020) may modulate the sensory input to the spinal cord as well as to supraspinal structures by preventing conductance failure and facilitating synaptic actions of sensory fibres. However, the effects of epidural polarization are long-lasting, for tens of minutes to hours rather than less than a second which is the case of PAD-related changes (Eccles *et al*., 1962; Rudomin & Schmidt, 1999; Rudomin, 2009). Changes evoked by epidural polarization may thus represent a particular form of modulation of input to the spinal cord.

With respect to the mechanisms behind the sustained increases in fibre excitability, morphological and molecular studies by D. Bennett’s group have demonstrated that Na^+^ channels within the proximal branching regions of afferent fibres are flanked by extra-synaptic GABA membrane receptors and that the excitability of these fibres is increased by GABA both in vitro (Lucas-Osma *et al*., 2018; Hari *et al*., 2022) and in vivo (Bradesi, 2010; Hari *et al*., 2022; Metz *et al*., 2023; Mahrous & Gonzalez-Martinez JA, 2024). The most recent findings of Bennett and Hari have also revealed that a short-lasting depolarization of these afferent fibres results in plateau potentials in the afferents that depend on both GABA and the depolarization (Krishnapriya Hari and David Bennett, personal communication, in preparation for publication).

The present study focused on the effects of GABA on afferent fibres traversing the dorsal columns in deeply anaesthetized rats in vivo in which changes in compound action potentials evoked in hindlimb peripheral nerves by dorsal columns stimulation were used as a measure of the excitability of afferent fibres. In this preparation, both GABA_A_ and the GABA agonist muscimol were shown to enhance the long-lasting increase in the excitability of afferent fibres following epidural polarization while two GABA antagonists (L655708, which blocks the extrasynaptic _α5_GABA_A_ receptors, and the non-selective GABA_A_ blocker bicuculine) had the opposite effect, decreasing the degree of post-polarization excitability and reducing the duration of the effects (Li *et al*., 2020). The long-lasting effects of local epidural polarization have been linked to the branching regions of afferent fibres at their entry to the spinal grey matter and to actions of GABA within these regions (Li *et al*., 2020). In the present series of experiments, we have explored the possibility that the long lasting increase in the excitability of afferent fibres by epidurally applied direct current (DC) depends on the depolarization of terminal branches of GABAergic interneurons as well as of glial cells within the branching regions of the stimulated afferents. As GABA may be released by glial cells (Yoon, 2012; Christensen *et al*., 2018; Woo *et al*., 2018; Hanani & Verkhratsky, 2021) as well as by interneurons, both may contribute to an increased concentration of GABA within and just ventral to the dorsal columns, as outlined in Fig. 1 and 2, GABA acting primarily via volume transmission. Such indirect actions on afferent fibres might result in an altered level of excitability of electrically stimulated afferent fibres but would also be likely to be induced under natural conditions, preventing branch point failures and thereby playing a critical role in axonal plasticity.

## Methods

All experiments were approved by the Regional Ethics Committee for Animal Research (Göteborgs Djurförsöksetiska Nämnd, permit no 5.8.18-16183/2019) and followed EU and NIH guidelines for animal care. The animals (26 female adult Sprague Dawley rats, 180-320 g, Janvier Labs, France) were housed under veterinary supervision at the Laboratory of Experimental Biomedicine at Sahlgrenska Academy with food and water ad libitum. Measures were taken to minimize animal discomfort by preceding the initial subcutaneous injections with a short period of isofluorane inhalation anaesthesia when the animal was unrestrained in a box. The number of animals was minimized by using protocols that allowed more than one question to be addressed in each experiment.

### Preparation

Anesthesia was induced with isoflurane (Baxter Medical AB, Kista, Sweden; 4% in air) followed by intraperitoneal administration of pentobarbital sodium (Apoteksbolaget, Göteborg, Sweden; 30 mg/kg) together with α-chloralose (Acros Organics, Geel, Belgium, 30 mg/kg).

The anaesthesia was supplemented at 3-4 hours intervals with an intraperitoneal injection of α-chloralose (3 mg/kg, up to 40 mg/kg). After initial surgical procedures and only when a deep level of anaesthesia had been established, the neuromuscular transmission was blocked by gallamine triethiodide (Sigma Aldrich, G8134) injected intravenously (via the tail vein) at an initial dose of 10 mg/kg supplemented with 5 mg/kg when needed and artificial ventilation applied via a tracheal tube by a respiratory pump (CWE; model SAR-830/P) set to maintain the end-tidal CO_2_ level at ∼3–4%. The core body temperature was kept at ∼38°C using servo-controlled heating lamps. To compensate for fluid loss, 10–20 ml of a glucose bicarbonate buffer were injected subcutaneously at the beginning of the experiments. The animal was monitored throughout the experiment with an ECG, via needle electrodes inserted subcutaneously over the thorax. The experiments were terminated by a lethal injection of pentobarbital resulting in ECG-verified cardiac arrest. In the hindlimb the peroneal (Per) and tibial (Tib) nerves were dissected, transected distally, and mounted on pairs of silver electrodes in a paraffin oil pool maintained at 32-35°C. The spinal cord was fixated with clamps, a laminectomy performed exposing the L1–L4 spinal segments and the cord covered with paraffin oil maintained at 35-37 °C. The dura remained intact apart from a small opening made for the drug-containing glass micropipette allowing it to touch the dorsal column (see below).

### Stimulation and epidural polarisation

Per and Tib group I muscle afferent fibres were stimulated at the level of Per and Tib motor nuclei where the most extensive branching of these afferents occurs. The location of these levels was guided by recording afferent volleys in group I afferent fibres in Per and Tib nerves rather than using the vertebrae as landmarks as the relationships between the L3-L4 spinal segments and the vertebrae may vary between individual animals (see Li *et al*., 2020 and diagrams in Nicolopoulos-Stournaras & Iles, 1983; Toossi *et al*., 2021 and Gerasimenko *et al*., 2019). The nerves were stimulated bipolarly using constant voltage stimuli at intensities 2–5 times threshold and the largest afferent volleys were evoked within the 2–3 mm length of the spinal cord (see Figure 1b in Li *et al*., 2020). The tungsten electrode was positioned on the dorsal column at the centre of this region. The dorsal column was stimulated monopolarly by a tungsten needle electrode insulated except for 20–30 μm at the tip (Microneurography active needle, UNA35FNM, FHC, Bowdoin, ME, USA; impedance 70– 400 kΩ) against a subcutaneous abdominal reference electrode. Single 0.2 ms constant current rectangular stimuli were delivered at intensities up to 60 μA. The stimuli were applied epidurally to avoid direct contact between the electrodes and the nervous tissue (see Holsheimer, 2002; Bikson *et al*., 2013; Holsheimer & Buitenweg, 2015; Jackson *et al*., 2017) but the layer of the cerebrospinal fluid between the dura mater and the dorsal columns was reduced by either indenting the intact dura or by draining the cerebrospinal fluid via small openings made in the dura to allow the drug containing micropipette to touch the cord surface.

Epidural polarisation was passed via the same tungsten electrode, based on having previously failed to find interference with the stimulus current stimulation under these experimental conditions (Bączyk and Jankowska 2014). The polarization was delivered via a custom-designed polarizer (Magnusson, D, Göteborg University). using the same parameters as in previous studies. The 1 μA depolarisation delivered for 1 min results in a maximal increase of the dorsal column fibre excitability within a radius estimated to be less than a millimetre and lasting for a post-polarisation period of minutes up to at least an hour (Jankowska *et al*., 2017). Changes in the size (area) of compound action potentials following epidural stimulation were used as a measure of the number of excited nerve fibres and thus of their excitability. Changes in the excitability were evaluated by comparing nerve volleys evoked by near-threshold stimuli (10– 30 μA) for a minimum of 5 minutes prior to passing DC during (1 min) and after DC application (for up to 30 minutes) using the same stimulus parameters The refractory period was evaluated by comparing responses to the second and the first of a pair of stimuli delivered in a series of standardized interstimulus intervals (0.55, 0.6, 0.65, 0.7, 0.75, 0.8, 0.85, 0.9, 1.0, 1.1 and 1.2 ms). The stimuli were delivered at twice threshold (2T) intensities (30-80 μA) allowing for a sharp onset and synchronisation of the nerve volleys but avoiding the spread of current over distances exceeding that of the epidural polarization. Changes in the absolute and relative refractory period were estimated by the ability and degree to which the fibres stimulated by the second stimulus elicited a nerve volley.

### Recording

The compound action potentials in Tib and Per nerves evoked by epidural stimulus used as a measure of the number of excited nerve fibres were recorded bipolarly by placing the transected nerves on a pair of silver-silver chloride hook electrodes 3-4 mm apart. The orthodromic afferent volleys evoked by peripheral nerve stimulation were recorded from the surface of the spinal cord close to the dorsal root entry zone by a silver ball electrode against a reference electrode inserted subcutaneously above the abdomen. Single records, as well as averages of 10 consecutive nerve volleys, were sampled and stored online (sampling frequency 33Hz; low passband filter set to 1 or 15 Hz and high-pass band filter at 5 or 3 Hz, resolution of 0,03 ms).

### Drug application

The drugs were allowed to diffuse from a glass micropipette (tip diameter 2.5-3 µm, R= 6-12 MΩ) in touch with the cord surface through a small opening in the dura about 0,5-1 mm rostral to the tungsten electrode. The following <u>solutions in 0.9% sodium chloride were</u> used: 200 mM GABA, Merck), 100 mM Gabazine, SR-95531 Merck) and 1 mM L-alpha-aminoadipic acid (L-AAA) A7275 Sigma. Unless stated otherwise, the diffusion continued during the whole period of recording to secure a relatively constant drug concentration. Effects following micropipette withdrawal were looked at only occasionally because the rate at which the drug was washed away could not be estimated.

### Analysis

The areas of nerve volleys were estimated using a custom-designed analysis programme (designed by E. Eide, Göteborg University, see Jankowska *et al*., 1997) and expressed in arbitrary units. The earliest components were measured within a time window of 0,4-0,7 ms from their onset (see boxed areas in Fig. 1 and 3), thereby excluding a non-linear summation with any later components and restricting the analysis to the fastest conducting fibres.

Furthermore, by verifying that the early components were evoked at latencies corresponding to those of afferent volleys in group I muscle afferents a contribution of group II afferents to the analysis was eliminated.

As the results from Per and Tib nerves were similar the data from both nerves were pooled together for statistical analysis unless stated otherwise. Student’s T-test was used to evaluate differences between normally distributed experimental data for each interstimulus interval and for selected records obtained during the post-polarization period.

### On the validity of the reported results

In order to decrease the number of experimental variables, the conditions under which the effects of DC were examined were greatly reduced; whenever possible these effects were tested under standard optimal conditions rather than a number of conditions. Thus, to eliminate the distance between the stimulation and polarization sites as a factor, the fibres were stimulated and polarised by the same electrode. To standardize the stimulus parameters the excitability and the refractory period of the fibres were routinely tested using near-threshold and twice-the-threshold stimuli.

The conditions under which GABA, gabazine and L-AAA were administered were less controllable. The choice to allow the drugs to diffuse from the micropipette in contact with the surface of the dorsal columns rather than using ionophoretic application was made to avoid the current needed to eject the drugs introducing an additional variable that would be difficult to control. However, the distance and velocity with which the drugs diffused and whether the diffusion rate differed for different compounds remained undefined. Ten minutes were, therefore, allowed for the diffusion of GABA and gabazine and 30 min for the diffusion of L-AAA over a distance of 1 mm from the stimulation site as fairly stable changes in the excitability of the tested fibres were usually detected after these periods.

When the drugs reduced the activation of only a proportion of dorsal column fibres, it could accordingly not be ascertained to which extent this might represent a failure of the diffused drug to block more than a limited number of nerve fibres or was due to the electrical stimuli targeting fibres within a larger radius than that over which the drugs diffused. Nevertheless, the possibility that the drugs failed to affect nerve fibres located deeper within the dorsal column does not appear to be the most likely explanation. This conclusion is drawn based on the findings that the effects were as potent on fibres stimulated over a longer distance (using twice threshold stimuli when analysing the refractory period) as on those located closer to the stimulation site targeted by near-threshold stimuli. In addition, while according to Holsheimer, 2002 and Holsheimer & Buitenweg, 2015 stimuli applied epidurally are effective only within a radius of a fraction of a millimetre, under our experimental conditions the drugs were likely to act within at least 1 mm, considering the distances between the tip of the micropipette containing the tested drugs and the epidural stimulation site.

Further unknown factors would include differences in the effects of both the current and the drugs used in this study on stem axons and on axon collaterals of afferent fibres traversing the dorsal columns, the branching regions of these fibres and, in particular, terminal branches of the GABAergic interneurons and glial cells. Future studies will thus be needed to define the specificity of changes evoked at the various sites of the proposed circuit.

## Results

### Polarization-evoked changes in the excitability of dorsal column fibres related to GABA

The modulation of long-lasting effects of epidural polarization in the branching region of afferent fibres was examined with respect to both the excitability and refractory period of the fastest conducting group I afferents from muscle nerves. These effects were examined during and following local administration of GABA and the GABA_A_ receptor blocker gabazine. To this end, GABA and gabazine were allowed to diffuse from a micropipette in contact with the surface of the spinal cord (Fig. 1B). The drug-containing micropipette was placed within less than a millimetre from the tip of the stimulating electrode, at a site where the largest afferent volleys were evoked within the L3-L4 lumbar segments by stimulation of the peroneal and/or tibial nerves (see Li *et al*., 2020), indicating that it overlaid the densest branching regions of afferents from these nerves as specified in Fig. 1A.

**Figure 1.**
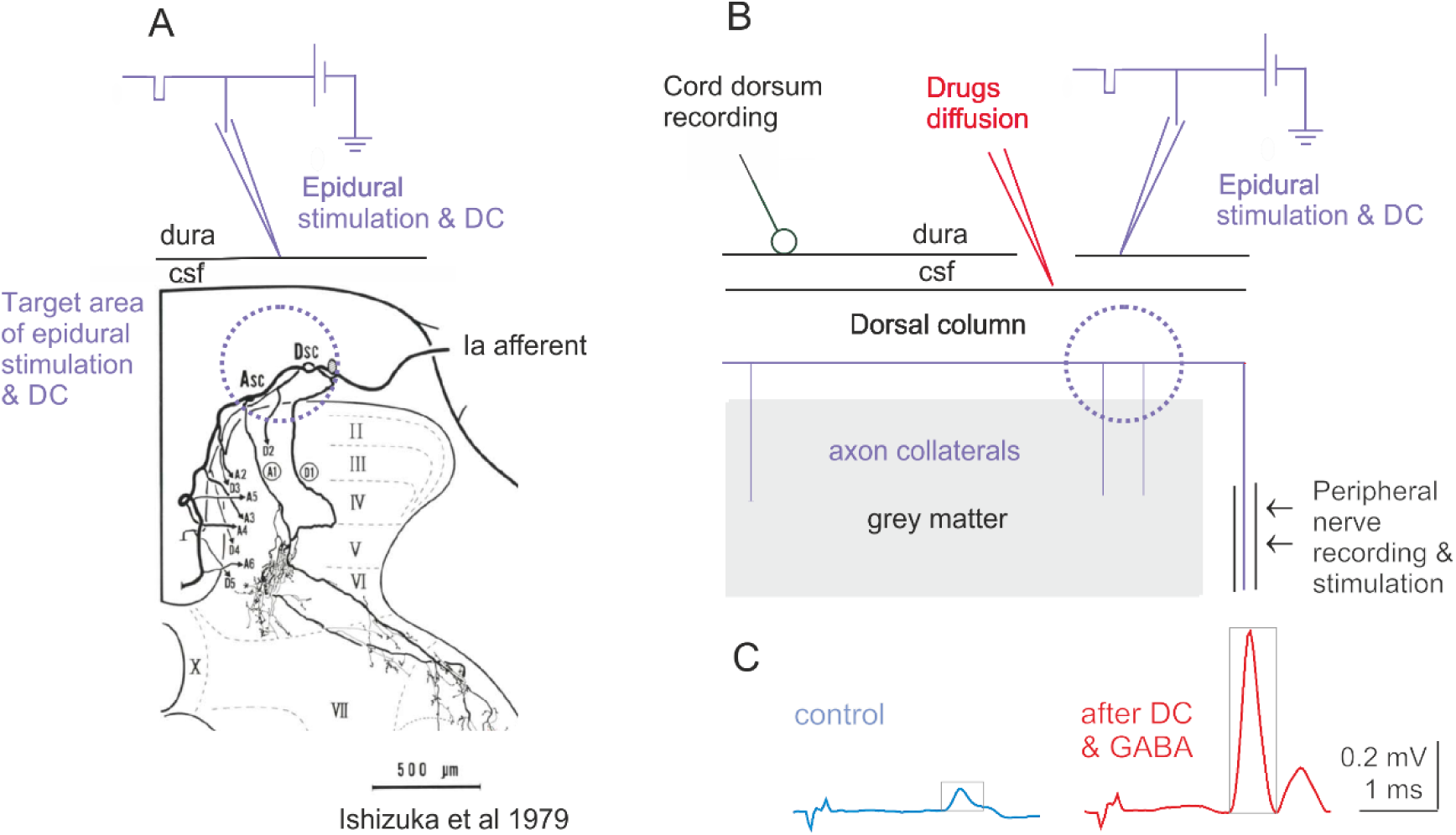
The experimental set-up. **A**, Drawing in the transverse plane illustrating the likely target area of the epidural stimulation Afferent fibres stimulated within the encircled area are represented by a fibre from the medial gastrocnemius, labelled and reconstructed by (Ishizuka *et al*., 1979). Modified from Fig. 1 in (Kaczmarek & Jankowska, 2018). The epidural tungsten electrode (purple) is used both to stimulate and polarize the fibres, **B,** Diagram of the stimulated fibres in a sagittal plane. The stem axons traversing the dorsal column (purple) are shown to give axon collaterals to the spinal grey matter. Modified from Fig. 2 in (Jankowska & Hammar, 2021). The region where the stimulation is most effective is encircled. Electrodes from right to left are: a tungsten electrode (purple) used both to stimulate and polarize the fibres, a glass micropipette (red) containing the drugs allowed to diffuse once in contact with the surface of the dorsal column via a small opening in the dura mater and a silver ball electrode (black) used to record afferent volleys from the peripheral nerves at the beginning of the experiment. Electrodes in contact with the transected peripheral nerves served both to stimulate the nerves to induce nerve volleys recorded from the cord dorsum and to record nerve volleys evoked by epidural stimulation. **C,** Examples of nerve volleys evoked by near-threshold epidural stimuli under control conditions (left) and the much larger volleys evoked by the same stimuli following epidurally applied DC (right). The size of the volleys was measured within the boxed areas.

**Figure 2.**
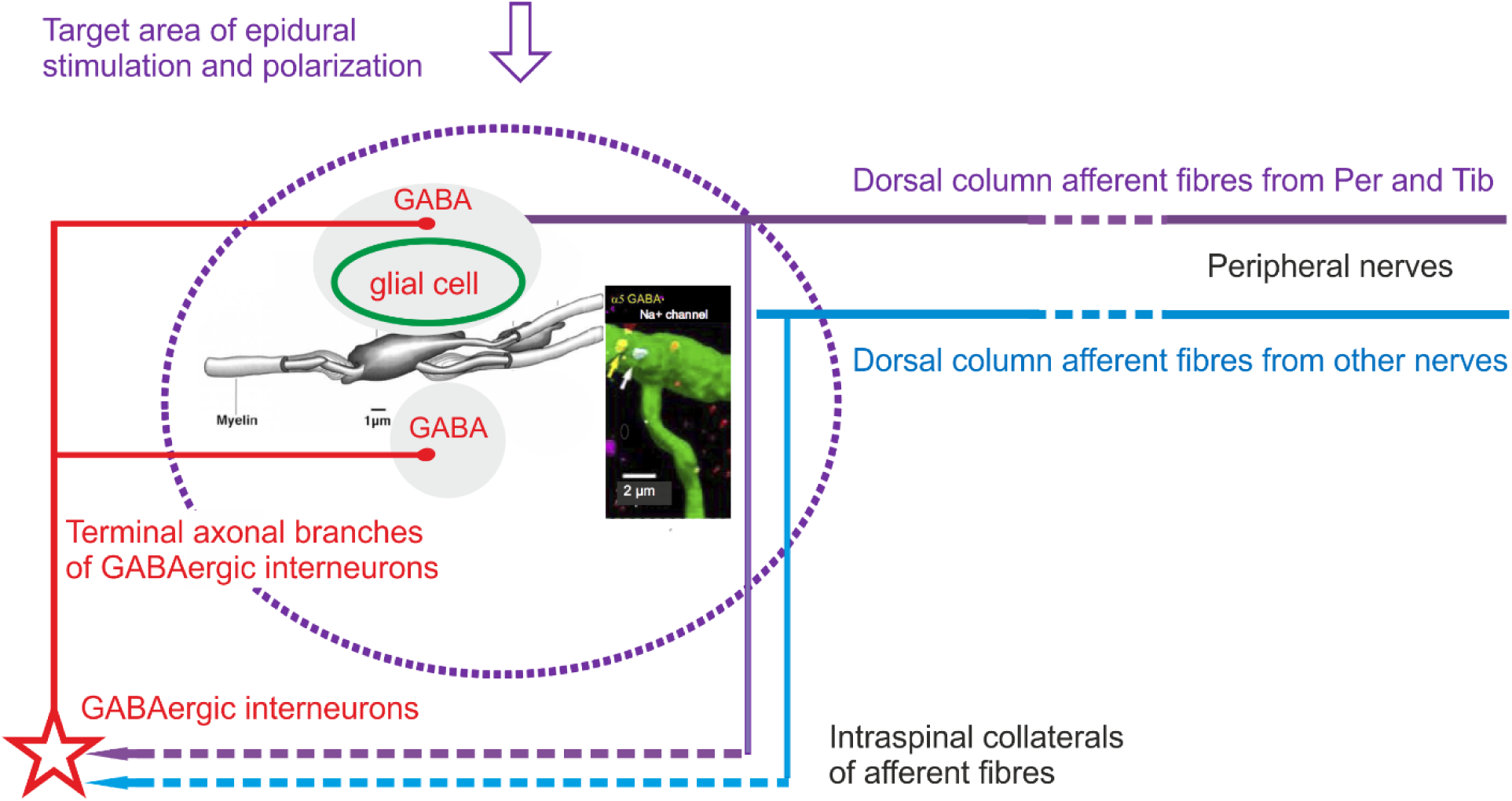
Hypothesized circuits of direct and potential indirect effects of epidural stimulation and epidurally applied DC including two potential sources of GABA released by epidurally applied DC. The area within which the epidurally applied DC is most effective is encircled. It includes the branching regions of group I muscle afferent fibres traversing the dorsal columns, illustrated with those described by (Nicol & Walmsley, 1991) and (Lucas-Osma *et al*., 2018) (left and right respectively). Shaded areas on either side of the branching points indicate areas within which GABA released by epidurally applied DC might affect the excitability and other properties of the afferent fibres. Two possible sources of GABA are indicated: the terminal branches of GABAergic interneurons (above and below; in red) and glial cells in the close vicinity (above; in green). Projections of epidurally stimulated afferent fibres to GABAergic interneurons located deeper within the spinal grey matter are indicated by dashed lines. For further explanations see text.

The neuronal targets of epidural stimulation and polarization and of the diffusing drugs within the encircled areas in Fig 1 A and B are specified in Fig. 2. These targets include sensory fibres traversing the dorsal columns (purple and blue) as well as terminal branches of GABAergic interneurons (red) and glial cells (green) close to branching regions of the afferent fibres. The figure also illustrates details of the morphology of the branching regions within and ventral to the dorsal columns as described by Nicol & Walmsley, 1991 and Lucas-Osma *et al*., 2018, including the area of several micrometres devoid of myelin sheet where extrasynaptic GABA receptors are found close to sodium channels.

The drugs diffusing from the pipette in contact with the surface of the spinal cord were found to reach the stimulated dorsal column fibres within a few minutes, as judged by the time elapsed until the nerve volleys recorded from the peripheral nerves showed signs of either decrease or increase (Fig. 3 B, D and F). However, the degree of these changes varied in individual experiments despite using the same concentration of the drug solutions, similar micropipette tip sizes and resistance and similar distances between the drug-containing micropipette tip and the stimulating tungsten electrode. The distances over which the diffusion occurred between the surface of the dorsal column and the depth at which the tested fibres traversed the dorsal columns and/or gave off axon-collaterals, might have varied, with considerabledifferences in the medio-laterally trajectory of the fibres at different rostrocaudal levels as well as for the particular subset of stimulated fibres. For these reasons, while the reported results show that the locally applied drugs affect the excitability of epidurally stimulated fibres, the degree of their effects can only be approximately estimated.

The decrease in the nerve volleys induced by epidural stimulation in the presence of gabazine (to about 80%; Fig. 3CD) in contrast to their increase by GABA (to about 120%; Fig. 3AB) indicates that the proportion of excited afferent fibres was decreased or increased respectively due to a decreased or increased excitability of these fibres. The decrease in the presence of gabazine also indicates the possibility that these fibres are under constant tonic facilitatory actions of GABA that may be counteracted by gabazine (Hari et al. 2022). The duration of post-application effects of GABA was not investigated as we could not control the rate with which it was washed away. Occasional observations indicated that the excitability of fibres remained increased for 15 min after the pipette containing GABA had been withdrawn (in 2 experiments) or after GABA ionophoresis had been terminated (Li *et al.,* 2020).

**Figure 3.**
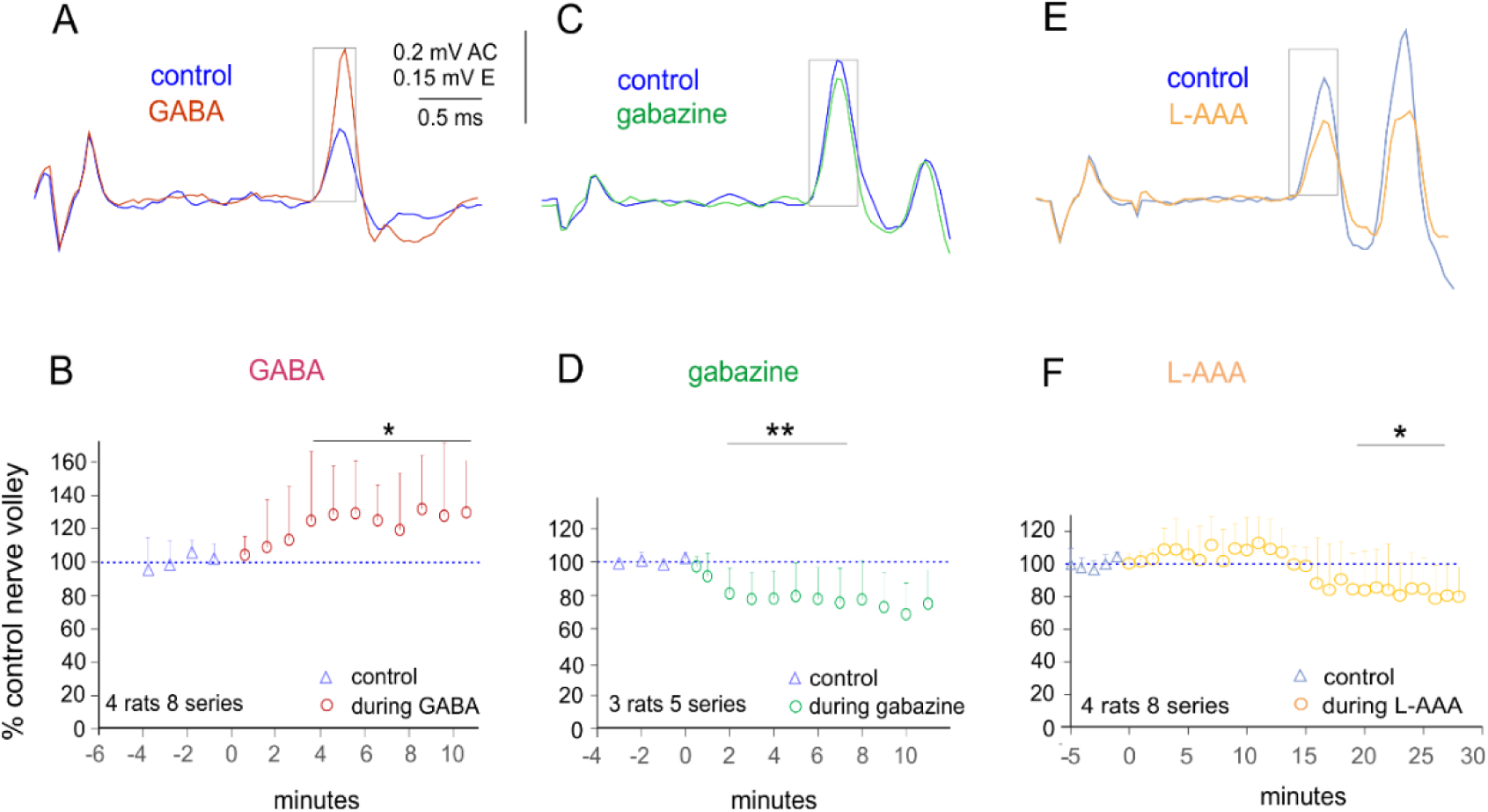
Effects of drugs modulating the excitability of dorsal column fibres. **A**, Examples of nerve volleys in the peroneal nerves evoked by epidural stimulation before (blue) and during the first 10 min of administration of GABA (red). Averages of 10 single records. **B**, Changes in the areas of the nerve volleys (ordinate) with respect to the mean areas before (blue) and during diffusion of GABA (red). The areas of the early components were measured in arbitrary units within the indicated windows. **C** and **D,** as in A and B but for nerve volleys recorded before (blue) and during diffusion of gabazine (green). **E** and **F**, as in **A** and **B** but for nerve volleys recorded before (blue) and during diffusion of astrocyte toxin L-alpha-aminoadipate, L-AAA (orange). Note similar effects of gabazine and L-AAA. * and ** indicate statistically significant changes when compared to the last control values (* p<0.01 - 0.05. ** p<0.001, Student’s t-test for paired samples).

Nerve volleys evoked by stimulation of the dorsal columns are severalfold increased by epidural polarization. As shown previously, DC (1µA for 1 min) applied to the surface of the spinal cord may increase these volleys not only during but also for minutes or even hours following the local depolarization of the dorsal columns, indicating that DC evokes a sustained increase in the excitability of dorsal column fibres (Jankowska *et al*., 2017; Bączyk & Jankowska, 2018). In the presence of GABA the sustained post-polarization increase in excitability was similar, irrespective of whether GABA was administered ionophoretically within the stimulated dorsal column (Li *et al*., 2020) or by diffusion from the surface of the spinal cord (present study). Fig. 4A shows that during diffusion of GABA the same stimuli evoked nerve volleys almost of the same size after (grey) as during (black) epidural polarization and that they were much larger than under control conditions (red). When effects of DC were tested during GABA diffusion, the nerve volleys decreased somewhat during the first 1-2 min of the postpolarization period (Fig. 4B) although in other experiments they were unchanged and did not show a decrease during the remaining tens of minutes. In contrast, when GABA effects were blocked by gabazine, the effects of epidural polarization declined to the pre-polarization level within the first 2-3 min (Fig. 4CD). These results are in line with those previously demonstrated when ionophoretically or systemically applied GABA antagonists (bicuculine and L655708 considerably reduced both the degree and duration of postpolarization changes in excitability (Li *et al*., 2020).

Taken together, these results demonstrate that GABA is needed for the long-lasting effects of DC. However, the considerable sustained post-polarization increase in excitability was previously found not only in preparations in which GABA was administered, either topically or ionophoretically but also in untreated preparations (Li *et al*., 2020). Hence under our experimental conditions, the background level of GABA appears to be sufficient for maintaining the post-polarisation effects on afferent fibres.

**Figure 4.**
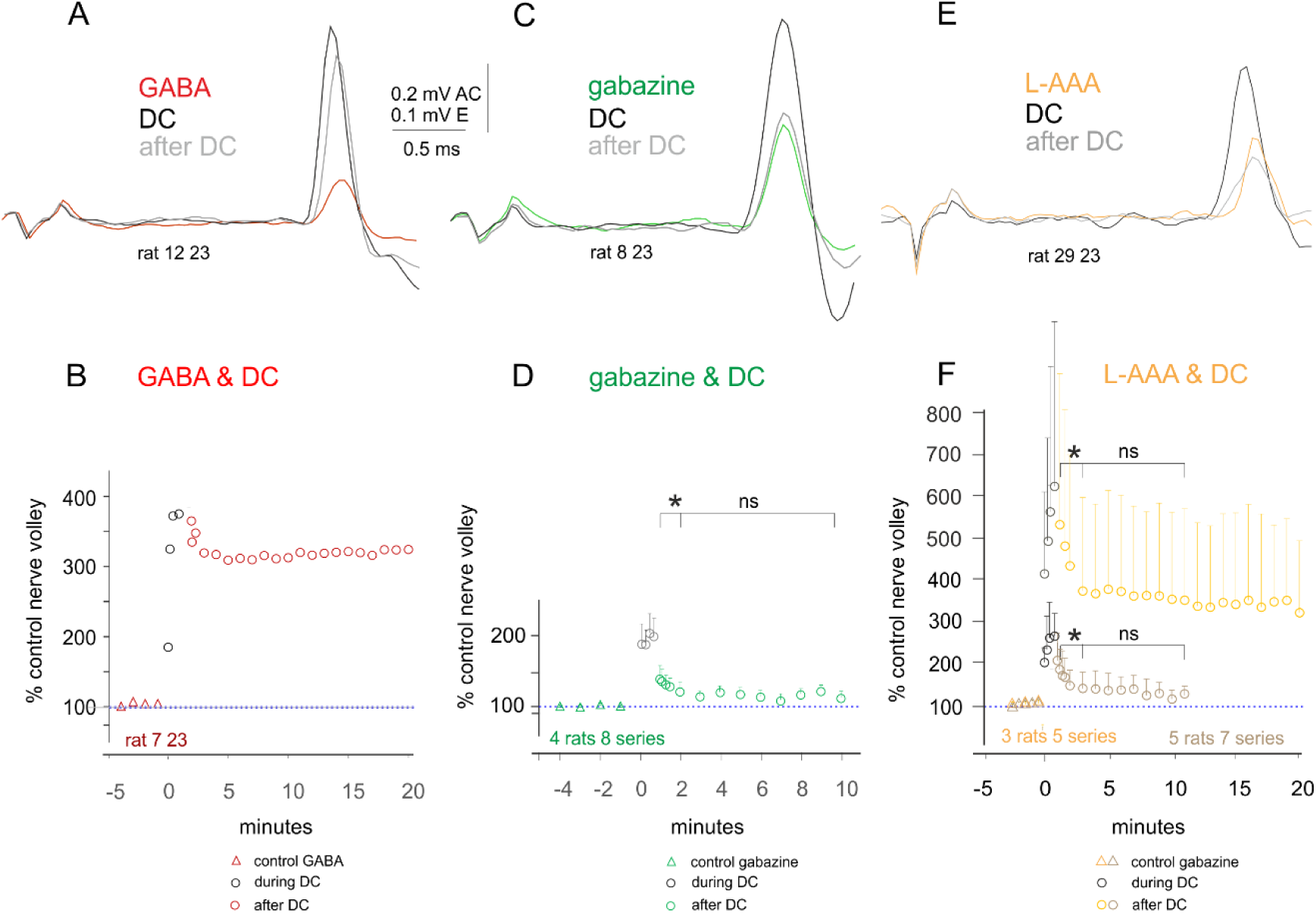
Effects of epidural polarization on the excitability of dorsal column fibres in the presence of GABA, gabazine and L-AAA. **Upper row**, Examples of nerve volleys in a peripheral nerve evoked by epidural stimulation before, during and following epidural polarization (1µA, 1 min) >30 min after the start of ongoing local diffusion of GABA (red), the GABA channel blocker gabazine (green) and the astrocyte toxin L-AAA (orange). Averages of 10 single records. Note the return of nerve volleys recorded during the post-polarization period to the control size in the presence of gabazine and L-AAA in **C** and **E** but not in **A**. **Lower row,** Changes in the areas of early components of the nerve volleys with respect to the mean areas of the records preceding DC (triangles), during DC (black circles) and following DC application (coloured circles); during diffusion of GABA (red), gabazine (green) and L-AAA (orange and brown in two samples; in 3 rats, five series and 5 rats, 7 series respectively). * indicates statistically significant differences between nerve volleys evoked immediately after the termination of DC (averages of 10 first volleys evoked at 1 Hz (Students t-test for paired samples). No differences were found between nerve volleys evoked at 3 and 10 min of the post-polarization period. However, the volleys evoked after the first 3 min levelled when they reached different sizes.

**Figure 5.**
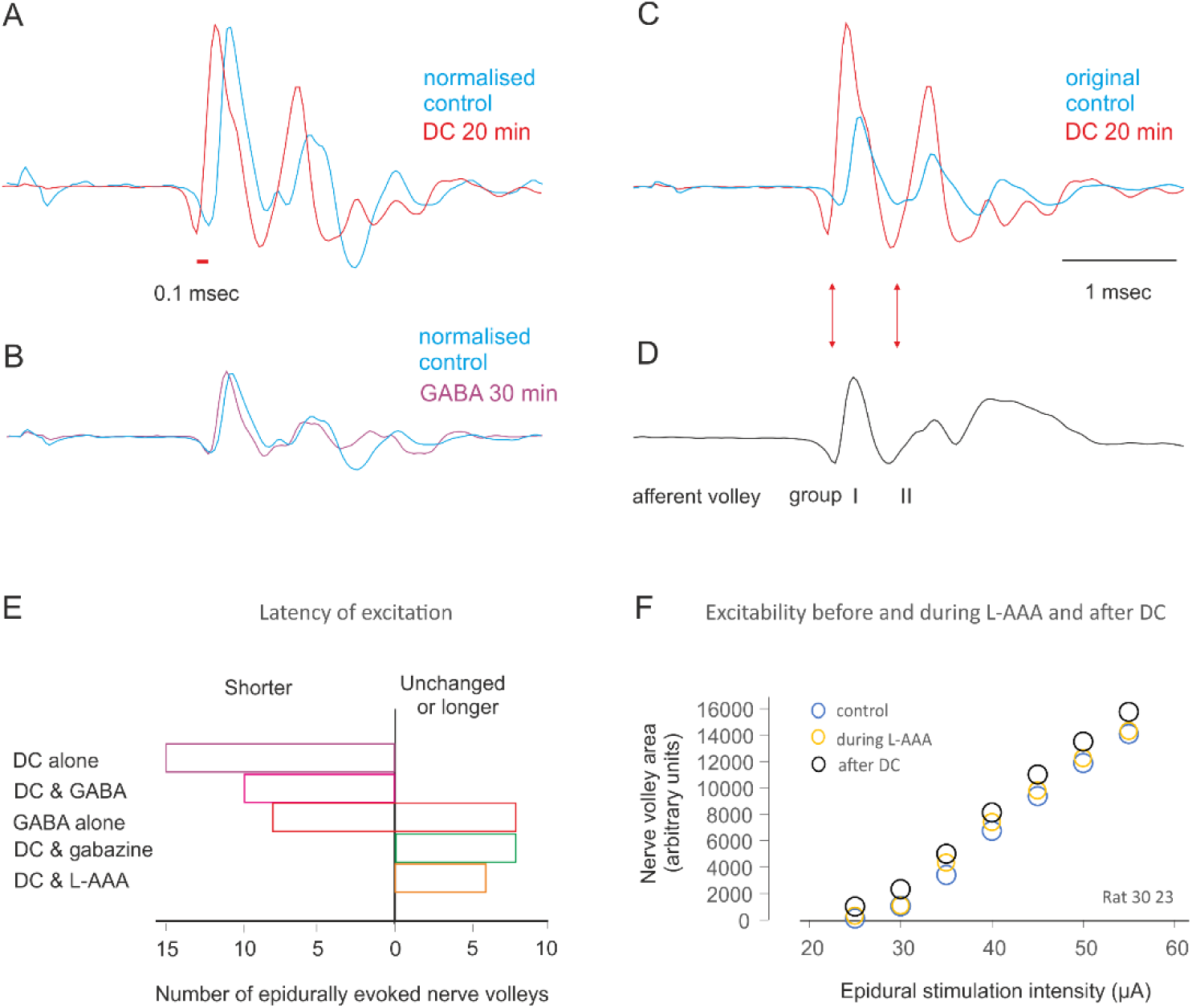
Effects of epidural polarization on the excitability of dorsal column fibres in the presence of GABA, gabazine and L-AAA estimated from the timing of nerve volleys induced by epidural stimulation. **A** and **B**, Examples of nerve volleys evoked by near-threshold stimuli before (blue) and following epidural polarization or in the presence of GABA (red). Amplitudes of the two volleys in a pair were normalized to compare their timing. **C**, Original sizes of similarly overlaid nerve volleys evoked in Per before and following epidural polarization, **D**, afferent volleys evoked by Per stimulation at 5T, recorded from the cord dorsum at the site of the epidural stimulation. **E**, Histograms of the latencies of nerve volleys shorter (as in A) or either unchanged or longer during the post-polarization period when DC was applied alone or in different combinations with concomitant diffusion of GABA, gabazine and L-AAA, as indicated. **F**, Comparison of the excitability of dorsal column fibres before and after DC applied in the presence of L-AAA. Data from an experiment in which the areas of the nerve volleys evoked by a series of epidural stimuli at the same intensity were not changed in the presence of L-AAA and only marginally increased during the post-polarization period. As the ranges of the stimuli used in such experiments varied because of differences in the threshold stimuli, the comparison could not be done for the population data but the relationships in F were representative for all 7 series in 5 rats.

The question of whether once the long-lasting effects of DC are induced, their maintenance depends on an elevated concentration of GABA within the branching region of the afferent fibres, i. e. whether they require the continuous actions of GABA on the fibres, has been addressed by Li *et al*., 2020. Their results demonstrate that the DC-evoked increase in the excitability was not significantly reduced when GABA channel blockers were applied during the post-polarization period (Fig 3E in Li et al. 2020) suggesting that GABA is primarily needed for the induction of the sustained effects.

The long-lasting increase in the excitability of nerve fibres during the post-polarization period was expressed not only in the size (area) of nerve volleys evoked by epidural stimulation but also in the timing of these volleys. As illustrated in Fig 1, 4 and 5, following epidural polarization the nerve volleys were not only larger but were also evoked at a shorter latency. When the size of nerve volleys evoked before and after epidural polarization was normalized, the rising phase and the peak of these volleys were shifted to the left (Fig. 5A); the shift was of the order of 0.03-0.18 ms and was present during at least 5-10 min of the postpolarization period. Such a post-polarization shift was found when the nerve volleys were evoked by 1.1-1.5 times threshold but not by stronger stimuli (Fig. 5E left), i.e. within the radius within which the excitability of many fibres was within the subthreshold fringe range. As no leftward shift occurred when nerve volleys were evoked by 2T or stronger stimuli, this phenomenon should be of minor consequence for effects of stimuli exceeding 2T, including changes in the refractory period of fibres described below. A similar, although smaller shift was found in about -half of nerve volleys evoked during the postpolarization period in the presence of GABA (Fig. 5B) but in none in the presence of gabazine blocking GABA receptors or L-AAA preventing the release of GABA (Fig 5E right).

A leftward shift characterised both the earliest and the later components of nerve volleys evoked by epidural stimulation. By relating these components to the afferent volleys evoked by stimulation of a peripheral nerve (Fig. 5 D) the earlier and later components may be attributed to compound action potentials evoked in group I and group II muscle afferent fibres respectively and indicate similar effects of DC on these fibres.

### Polarization - evoked changes in the refractory period of dorsal column fibres related to GABA

GABA-related changes in the excitability of afferent fibres were matched by changes in the refractory period of epidurally stimulated fibres. When two stimuli are applied to a nerve or a nerve tract, and the interstimulus intervals are decreased within the relative refractory period, the compound action potentials evoked by the second stimulus gradually decrease, reflecting the reduced number of fibres that remain excitable at the respective interval until no fibres can be re-excited within the absolute refractory period. In the case of epidurally evoked nerve volleys, those following the 2^nd^ stimulus overlapped with the late components of the volleys evoked by the 1^st^ stimulus. The measurements of their area therefore required that the second volleysare subtracted from the sum of the first and second volleys, as illustrated in Fig. 6 A & B. Records in Fig. 6B show that in the presence of GABA considerably larger responses were evoked at all interstimulus intervals corresponding to the relative refractory period. They also show that at the two shortest intervals the responses were present during GABA administration while they were absent at control conditions. This indicates a shortening of the absolute refractory period by about 0.2 ms. Such a shortening has previously been demonstrated following epidural depolarization (Fig. 6C, see Jankowska *et al*., 2022) and the present series of experiments shows that it may also be elicited following locally applied GABA (Fig 6 D).

**Figure 6.**
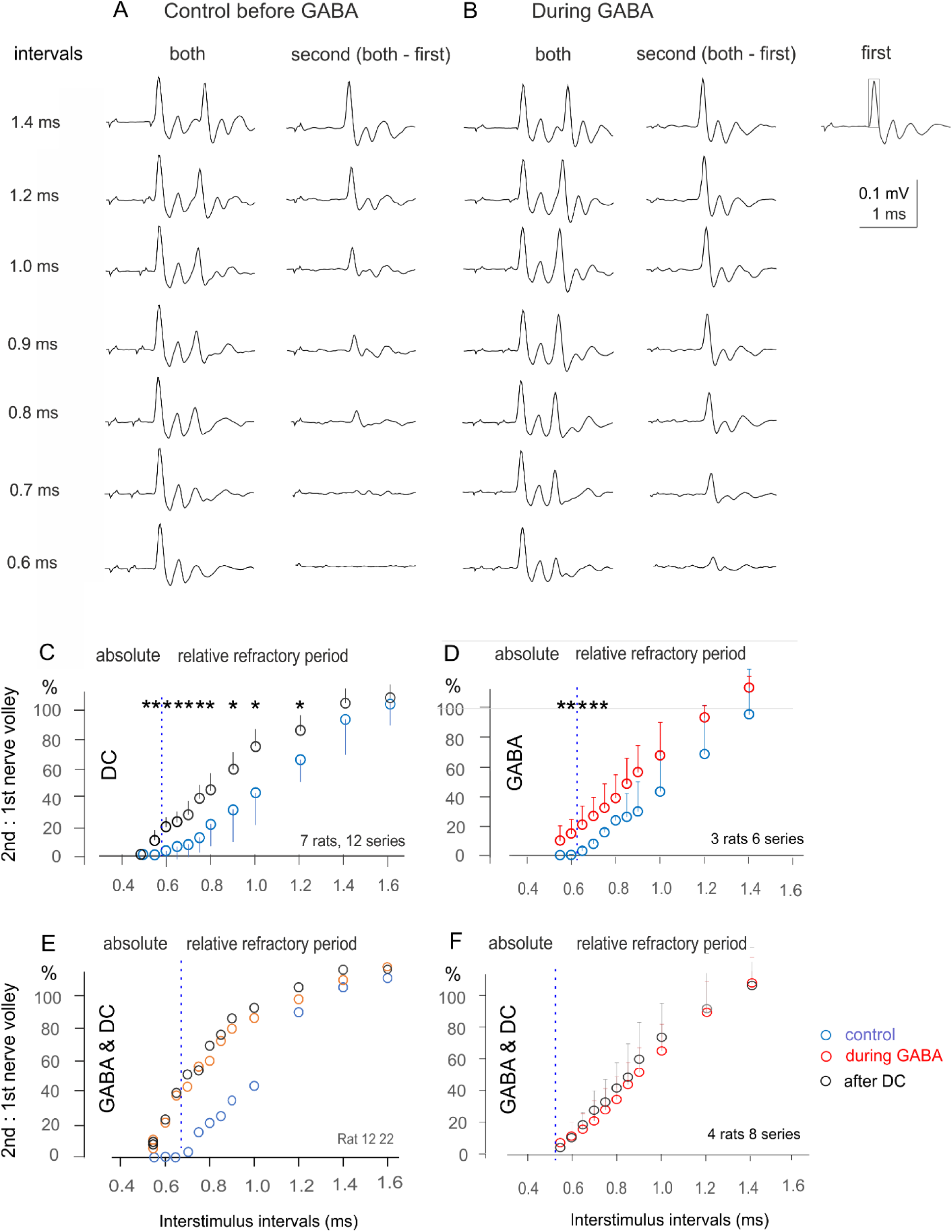
GABA and DC have similar effects on the refractory period of afferent fibres in the dorsal columns. **A** and **B**, Examples of nerve volleys evoked by pairs of epidural stimuli at decreasing interstimulus intervals under control conditions (A) and during diffusion of GABA (B) respectively. The records in the left columns show responses evoked by the second stimulus overlapping with late components of those evoked by the first stimulus. The nerve volleys evoked by the first stimulus were subtracted from the joint effects of the two stimuli (records to the right) allowing a comparison of volleys preceded and not preceded by the first stimulus. The comparison involved the areas of the early components (boxed in B, third column). **C**, The area of the nerve volleys evoked by the second stimulus related to those evoked by the first stimulus at decreasing interstimulus intervals under control conditions (blue) and following epidural polarization (black). (replotted from Fig. 2 in Jankowska *et al*., 2022). The minimal relative refractory periods under the two conditions are indicated by the vertical dotted lines. The shortening of the refractory period during the post-polarization period was associated with the leftward shift of the plots. **D,** refractory period of nerve volleys evoked before (blue) and during (red) local application of GABA (15-20 min after the diffusion commenced) with statistically significant differences for intervals 0.55 ms (p=0.031), 0.6, 0.65, 0.7 ms (p=0.013) and 0.75 ms (p=0.031). Student’s t-test for paired samples. Note the similarity between the curves in C and in D. **E**, Data from one of the experiments in which the refractory period was shortened by GABA but remained unchanged following the subsequent epidural polarization. **F,** Mean changes in the refractory period following GABA diffusion (red) and when GABA diffusion was combined with epidural polarization (dark grey) in 4 rats (8 series in which no statistically significant differences (Student’s t-test) were found between nerve volleys recorded during GABA application before and following DC at any interstimulus intervals).

**Figure 7.**
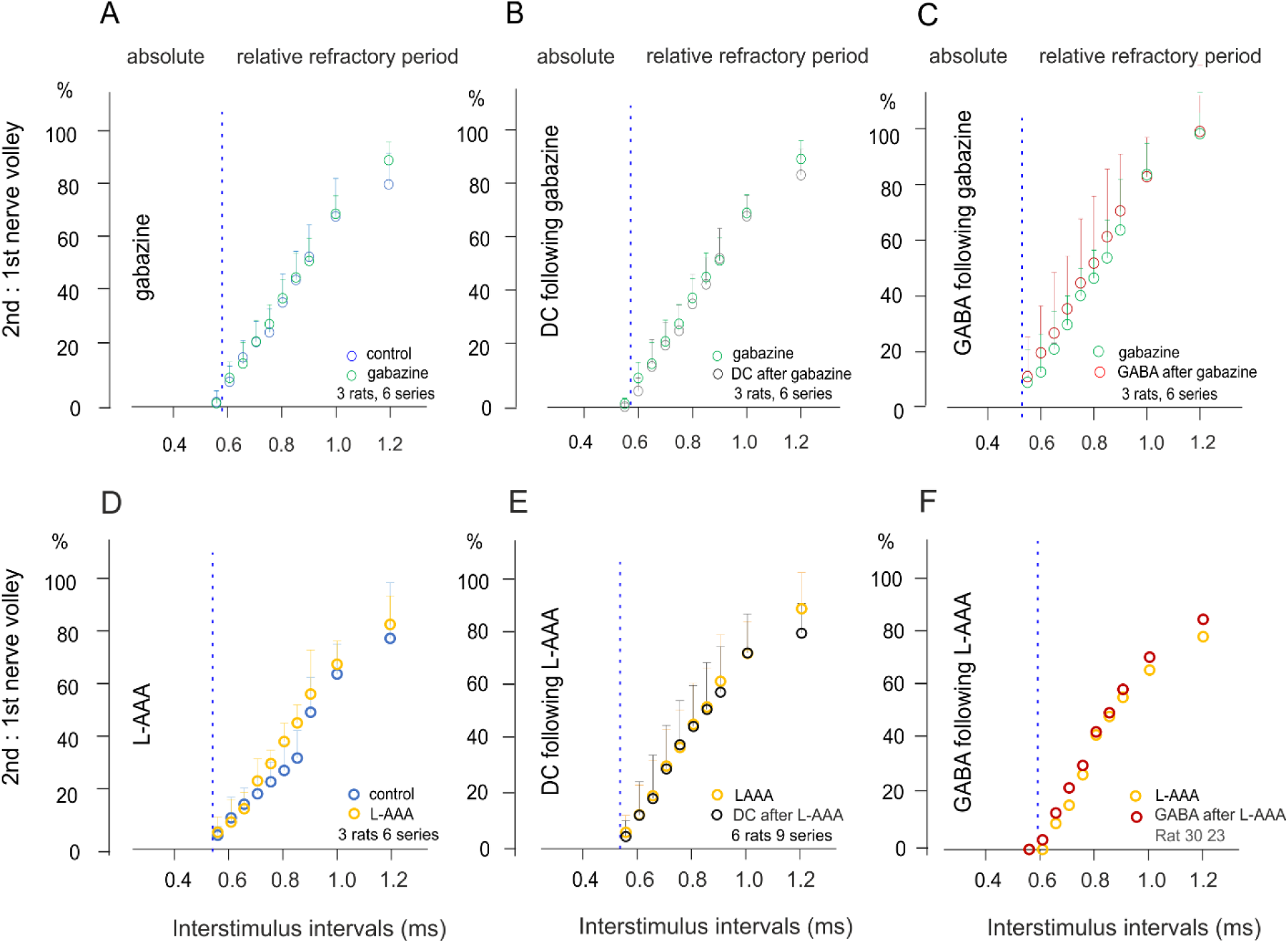
Effects of the GABA receptor channel blocker gabazine and the glial toxin L-AAA on the refractory period of dorsal column fibres. **A**, Refractory period under control conditions (blue) and following gabazine (green; 3 rats 6 series) administration. B, Comparison of the refractory period during administration of gabazine (green) and following the subsequently applied epidural polarization (black 3 rats, 6 series). **C**, As in B but comparing the refractory period during the administration of gabazine (green) and the subsequent administration of GABA (red; 5 rats, 10 series). D, Refractory period under control conditions (blue) and following glial toxin L-AAA (orange; 3 rats 6 series). E, Comparison of the refractory during administration of L-AAA (orange) and following the subsequent epidural polarization (black; 6 rats, 9 series). **F**, As in **E** but comparing refractory period during administration of L-AAA (orange) followed by GABA (1 series). Similar marginal differences were observed in two additional series. No statistically significant differences were indicated by Student’s t-test between data sets in A-F.

Once the GABA-evoked shortening of the refractory period had been established, epidural polarization failed to induce any further shortening, irrespective of whether the individual or mean sizes of nerve volleys evoked by epidural stimulation at different interstimulus intervals were compared (Fig. 6 E and F).

These effects indicate that not only the increased excitability but also the shortening of the refractory period of afferent fibres may be attributed to a DC-induced increase in the release of GABA and the resulting change in its concentration within the branching region of these fibres. They are also in keeping with previous observations that GABA has none or only minor additional effects on the sustained increase in the excitability evoked by DC (see above and Li *et al*., 2020) when DC effects are maximal.

In order to estimate the degree to which effects of epidural polarization on the refractory period might be secondary to GABA released by DC, GABA-related changes were examined not only when epidural polarization was applied together with GABA but also when GABA effects were blocked by gabazine.

Gabazine administered by diffusion had only a weak effect on the refractory period by itself (Fig 7A), However, it effectively prevented the shortening of the refractory period both by epidural polarization (Fig. 7B) and by GABA (Fig. 7C). The effects of gabazine are thus in keeping with the DC-evoked shortening of the refractory period being secondary to the DC related release of GABA.

### Sources of GABA released by epidural polarisation

As indicated in the introduction and outlined in Fig. 2, GABA acting on branching regions of afferent fibres could be released not only by GABAergic interneurons but also by glial cells Some GABAergic interneurons were found to project to the areas within as well as ventral to the dorsal columns (David Bennett, personal communication; Hari *et al.,* 2022) and epidurally applied DC could result in an increased amount of GABA released by these neurons by depolarizing their terminal branches (see Discussion). With respect to the glial cells, it is unknown which types of glial cells would be involved and whether such cells have the same properties as glial cells affecting neurons and synaptic transmission and whether they express GABA (as 20% of the dorsal horn astrocytes in the turtle spinal dorsal horn according to Christensen et al 2018). Provided they are GABAergic and located within the region within which axon collaterals are given off by dorsal column fibres, they could constitute another source of GABA released by epidural polarization.

Given that GABA released by glial cells and the GABAergic interneurons would be similarly counteracted by gabazine, it greatly narrows the possibility to differentiate between effects of GABA from these two sources. In two series of experiments, we nevertheless verified the potential contribution of GABAergic astrocytes to the DC effects. To this end, we examined whether the astrocyte toxin L-AAA (Pereira *et al*., 2021; Xu *et al*., 2021) would counteract the increase in the excitability and the decrease in the refractory period of epidurally stimulated afferent fibres. We also examined whether GABA-expressing astrocytes are present close to the branching regions of afferent fibres (Emelie Joelson and Ingela Hammar, in preparation).

Fig 3F shows that the excitability of epidurally stimulated nerve fibres was reduced in the presence of L-AAA. The nerve volleys evoked by epidural stimulation began to decrease within about 15 min from the onset of the diffusion of L-AAA but the decrease became statistically significant only after about 20 min. The effects of L-AAA thus resembled the effects of gabazine, although they were weaker and induced with a longer delay. The overall decrease in the excitability of epidurally stimulated nerve fibres by L-AAA was nevertheless marginal as indicated by the comparison of the effectiveness of stimuli of successively higher intensities before and in the presence of L-AAA. This is illustrated in Fig.5F with the data from one of the experiments because differences in the ranges of effective stimuli in various animals precluded pooling the data. However, similar relationships were found in all 7 series of records in 5 experiments. L-AAA also had only marginal effects on the refractory period of epidurally stimulated fibres (Fig. 7D).

In contrast to these weak effects, L-AAA effectively counteracted the long-lasting effects of epidural polarization albeit depending on how strong the effects of DC were during its administration. When DC evoked a moderate increase in the excitability in the presence of L-AAA (lower plot in Fig, 4F), the post-polarization increases in the excitability were short-lasting, the excitability returning towards the control level within the first few minutes. When the DC evoked-increase in the excitability exceeded 300%, the decline was as rapid but levelled out at a higher level (upper plot in Fig, 4F). The shortening of the refractory period of epidurally stimulated afferent fibres induced by DC or GABA was likewise counteracted in the presence of L-AAA. As shown in Fig 7, nerve volleys of similar size were now evoked by the second stimulus at increasing interstimulus intervals before and following the epidural polarization (Fig. 7E) and GABA administration (Fig. 7F).

Taken together the similarities between the local effects of gabazine and L-AAA on long-lasting DC-evoked changes in the excitability of afferent fibres are compatible with a similar contribution of glial cells and GABAergic interneurons to GABA-attributable induction of effects of epidural polarization.

Whether glial cells are critical for maintaining the long-lasting DC effects once they have been induced was not tested. Such tests would require that L-AAA was applied during the plateau phase of DC evoked increase in the excitability. However, L-AAA had to diffuse for about 30 min before its effects were established, precluding a reliable quantification of these effects.

Nevertheless, in one experiment in which this was tested, the excitability of afferent fibres following L-AAA administration during the post-polarization period was found to be decreased by 20-30%.

### The effects of DC and GABA depend on the distance from the branching regions of the afferent fibres

The results presented in the preceding sections were obtained when afferent fibres were stimulated at the sites corresponding to the regions of the densest collateral branching of muscle spindle afferents (Li *et al*., 2020). The analysed fibres belonged to the fastest conducting afferent fibres in the peroneal and/or tibial nerves and the greatest numbers of axon collaterals of group Ia afferent fibres would be given off over the peroneal and tibial motor nuclei targeted by them.

The anterior tibial and extensor digitorum longus motor nuclei are located within a more narrow stretch of the spinal cord (in the L3 and the most rostral part of the L4 segment) than motor nuclei of the posterior tibial nerve, including those of the triceps surae, plantaris and flexor digitorum longus (extending over the L3-L5 segments) (see Fig. 5 in Nicolopoulos-Stournaras & Iles, 1983 and Toossi *et al*., 2021; see also Gerasimenko *et al*., 2009 and Calvert *et al*., 2019) but they overlap within 2-4 mm. In most experiments, the DC effects on Per and Tib afferent fibres were accordingly similar and were pooled together in Figs 1-3. Nevertheless, in some experiments the effects differed, as illustrated in Fig. 8 A and C.

**Figure 8.**
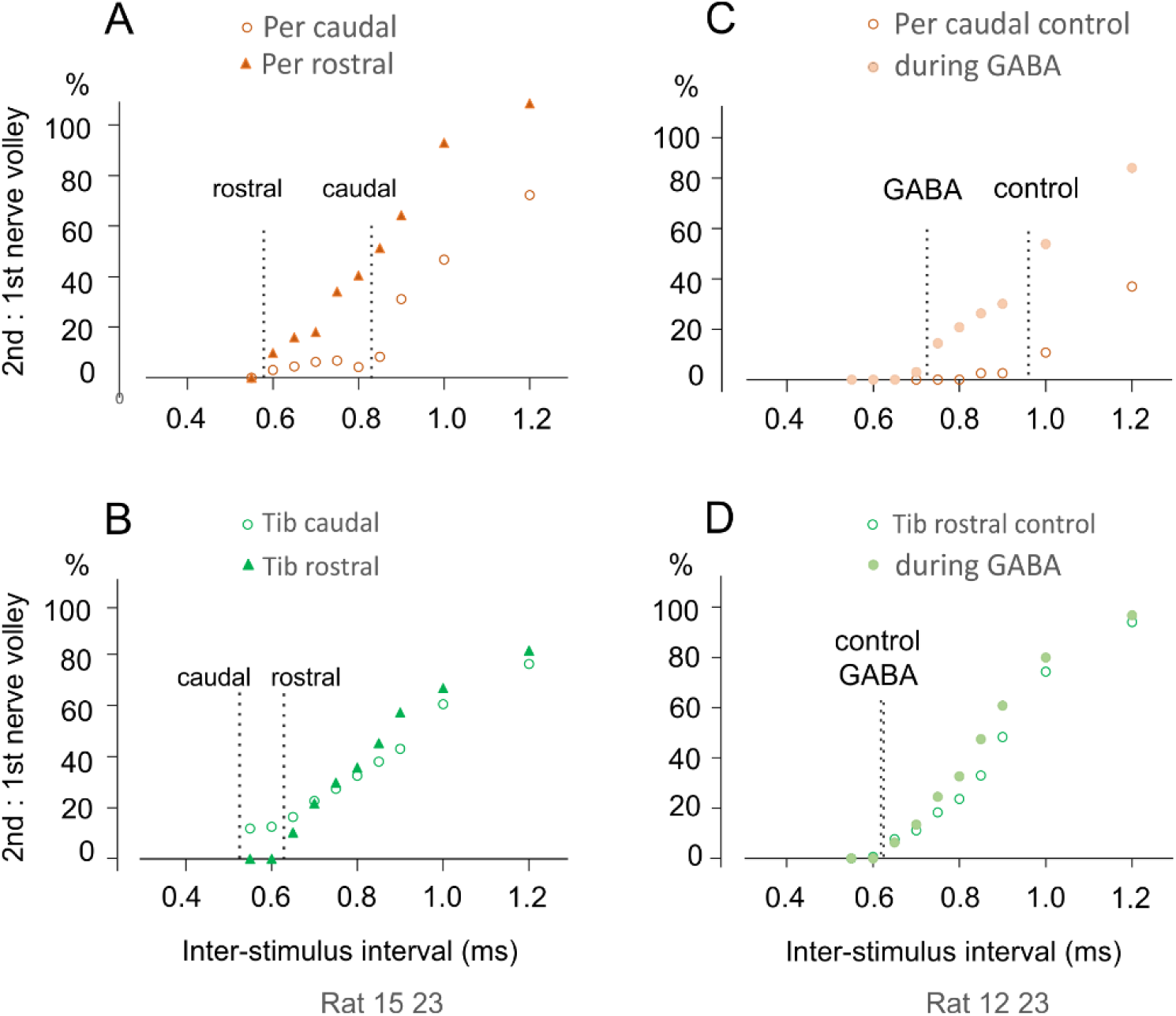
Differences in the ranges of the refractory period of Per and Tib afferent fibres stimulated within the rostral and caudal part of the lumbosacral enlargement. **Left and right panels**, data from two rats in which dorsal columns were stimulated at two different locations 6 mm part apart. A and B, Open circles and triangles are for nerve fibres stimulated more rostrally and caudally respectively. Note differences in the duration of the absolute refractory period of Per and Fib afferent fibres. **C** and **D,** Open and filled circles are for fibres stimulated before and during administration of GABA at a more rostral level for Per and a more caudal level for Tib. Note that GABA shortened the duration of the absolute refractory period in **C** but not in D. Vertical dotted lines indicate the borders between the absolute and relative refractory periods.

The ranges of the refractory period were compared at several rostrocaudal levels 2 mm apart. They were related to the levels at which maximal afferent volleys from group I afferents were recorded from the cord dorsum and at which maximal synaptically mediated components of these potentials were evoked from group II muscle afferents (see Fig 5D; see also Fig 1D and E in the recent analysis of cord dorsum potentials by Taccola *et al*., 2023.

When the dorsal columns were stimulated within the caudal part of the lumbosacral enlargement, the peroneal fibres failed to respond to the 2^nd^ stimulus when the interstimulus intervals were <0.7-0.9 ms (Fig. 8A and C, open circles) while the minimal inter-stimulus intervals at which fibres of the tibial nerve responded were 0.55-0.65 ms (Fig, 8B, open circles). When the dorsal columns in the same experiments were stimulated more rostrally, the ranges of the relative refractory periods of fibres in the Tib and Per nerves were similar, the fibres in both nerves responding to the 2nd stimuli at 0.55 -0.65 ms minimal inter-stimulus intervals or tended to be longer for the tibial fibres (Fig. 8B). No such differences were seen when the fibres were stimulated in peripheral nerves or in the dorsal roots for which the minimal interstimulus intervals are longer (Jankowska *et al*., 2022). These differences disappeared when the refractory period was shortened by GABA and/or DC (Fig. 8C and D).

One of the main reasons for the variability in the effects of DC and GABA in individual rats might thus be that the afferent fibres were stimulated at different distances from the regions with the densest collateralisation. It may nevertheless be noted that even when GABA_A_ receptors were blocked by gabazine, preventing the DC-evoked shortening of the refractory period, the range of the relative refractory period (0.55-1.2 ms) was as that found in control records (Fig. 6) and much shorter than the range of the refractory period characterising the unbranched compartments of afferent fibres in peripheral nerves (1.2-∼3 ms; see Jankowska *et al*., 2022).

## Discussion

The aim of this study was to examine how GABA combined with depolarization of afferent fibres at the level of their entry to the spinal grey matter modulates input to the spinal cord. The reported modulatory effects of GABA differ in several ways from those on synaptic transmission to spinal relay neurones that are closely linked to primary afferent depolarization (PAD). Firstly, at the level of the most proximal branching points of afferent fibres the effects of GABA are evoked by volume transmission, GABA being released at a distance, in contrast to being evoked by axo-axonic synapses formed with distal compartments of primary afferents. Secondly, the GABA-dependent DC effects are considerably longer-lasting, for tens of minutes or hours, rather than for less than a second for PAD evoked by single stimuli or short trains of stimuli. Thirdly, they may involve afferent fibres of several modalities rather than specific categories of afferents as is the case of PAD. Modulation of afferent input at the level of its entry to the spinal cord may thus play a different functional role and have a different impact on therapeutic procedures.

### Is the long-lasting increase in excitability of afferent fibres providing input to the spinal cord primarily due to the polarization of these fibres or secondary to GABA release and extrasynaptic GABA actions?

It has previously been established that the long-lasting increase in the excitability of afferent fibres induced by epidural polarization is enhanced by GABA and counteracted by GABA antagonists (Jankowska *et al*., 2017; Li *et al*., 2020). One of the main questions addressed in this study has been to what extent this effect depends on the *depolarization of afferent fibres themselves* or *on DC actions on preterminal branches of GABA-releasing interneurons or GABAergic glial cells*.

DC applied epidurally would affect nerve fibres within a certain radius but with the effectiveness rapidly declining within about 1mm distance (Holsheimer, 2002; Holsheimer & Buitenweg, 2015). However, as indicated in Fig. 2, nerve fibres located within this radius may include not only afferents from peripheral nerves but also terminal nerve branches of GABAergic interneurons projecting towards the dorsal columns. Epidural polarization might also affect glial cells surrounding the afferent fibres, constituting another potential source of GABA. Furthermore, the polarization might also facilitate the activation of GABAergic interneurons targeted by the epidurally stimulated afferent fibres.

Which of these would most readily take place may depend on several factors. Nevertheless, as terminal branches of nerve fibres are consistently excited at a lower threshold than their target cells (Baldissera *et al*., 1972; Gustafsson & Jankowska, 1976), the release of GABA from the terminal branches of GABAergic interneurons and glial cells might be affected at lower stimulus intensities than the afferent fibres in the dorsal columns. If so, the indirect effects of epidural polarization on the epidurally stimulated fibres might be as, or even more essential than the direct effects of DC, In order to estimate the relative contribution of GABA-related indirect effects of epidural polarisation, we compared the excitability and the refractory period of dorsal column fibres under three conditions: (i) during GABA diffusion not combined with DC (ii) when direct effects of DC on afferent fibres were evoked without the concomitant effects of GABA (GABA receptors having been blocked with gabazine) and (iii)_)_ when DC could affect the dorsal column fibres both directly and indirectly, by acting on any sources of GABA in their vicinity.

Effects of GABA application without fibre polarization were relatively moderate as shown in Figs. 3B, 6D, 8D, as were effects of DC evoked under conditions when effects of GABA were prevented by gabazine (Figs. 4D, 7B). In contrast, much stronger and longer-lasting effects were elicited following epidural polarization when it could affect dorsal column fibres both directly and indirectly by increasing the amount of GABA released close to the stimulated fibres (Figs. 4B, 6E).

Provided that the long-lasting increase in the excitability reflects a sustained depolarization of nerve fibres, its future analysis will be greatly assisted by the recent demonstration of plateau potentials in afferent fibres by David Bennett and Krishnapriya Hari (personal communication). With respect to the issues addressed in the present study, their finding that joint actions of depolarization and of GABA are necessary for the induction of these plateau potentials is particularly important. Neither depolarization, in particular intra-axonally evoked depolarization of single fibres on its own, nor the increased concentration of GABA in the extracellular space in their in vitro preparation were found to suffice for initiating the post-polarization plateau potentials.

We consider the reported effects of GABA to be due to its volume transmission (Agnati *et al*., 1995; Agnati & Fuxe, 2000b; Agnati, 2010; Gianni, 2023) in view of the demonstration of predominant or even exclusively extrasynaptic GABA membrane receptors within the proximal branching regions of afferent fibres, and because the extra-synaptic GABA_A_ receptor antagonist L655708 abolished the sustained effects of epidural polarization (Lucas-Osma *et al*., 2018; Hari *et al*., 2022). Within the distal compartments of afferent fibres the relative contribution of GABA modulatory actions evoked via extrasynaptic receptors and via axo-axonal synaptic contacts was concluded to be the opposite, the latter dominating (see in particular Fig. 3i and extended Fig.1 in Hari et al 2022). For the likely functional consequences of such different morphological substrates of effects of GABA within the distal and the proximal branching regions of the afferents see below.

### The relative contribution of effects of GABA released by GABAergic interneurons and by glial cells

The comparison of the effects of gabazine and L-AAA indicates striking similarities. The elimination of GABA released by glial cells (by L-AAA) and by GABAergic interneurons (by gabazine) might thus explain why L-AAA and gabazine counteract the long-lasting effects of epidural polarization similarly effectively, in support of the possibility that both glial cells and GABAergic interneurons are the source of GABA released by epidural polarization.

However, the properties of the main types of glial cells were reported to differ in several respects (see e.g. Bradesi, 2010; Serrano-Regal *et al*., 2020; Hanani & Verkhratsky, 2021) and only a proportion (20%) of astrocytes in the turtle spinal dorsal horn were found to express GABA (Christensen *et al*., 2018) and possibly operate as both “GABAceptive and GABAergic” cells (Goulding *et al*., 1991; Yoon, 2012; Woo *et al*., 2018) At the present stage of our knowledge we may thus only speculate about the contribution of glial cells as compared to the contribution of GABAergic interneurons. A relatively strong effect of GABA released by glial cells would for instance be favoured by a proximity between these cells and the branching regions of the afferents while the terminals of the GABAergic interneurons might be not numerous enough, nor close enough to these branching regions to initiate the sustained excitability changes by themselves.

This would be in keeping with the failure to evoke a sustained increase in the excitability of afferent fibres during the post-polarization period when the contribution of GABA from glial cells is eliminated by L-AAA, together with the weak increase in the excitability of afferent fibres evoked by GABA diffusion from the surface. As glial cells respond to changes in external field potentials, epidural polarization might affect not only the terminal branches of GABAergic interneurons but also glial cells, the effects on the latter being further enhanced by gap junctions between glial cells (see Ji *et al*., 2019), and the joint outcome more potent.

### Functional consequences of DC-evoked release of GABA acting via volume transmission

The DC-evoked increase in the excitability of afferent fibres traversing the dorsal columns may result in a considerable increase in the number of fibres activated by electrical stimuli, and thus improve the outcome of epidural stimulation. DC may also improve the propagation of nerve impulses along afferent fibres activated by natural stimuli by preventing branch point failures as well as reducing delays incurred at these points (for references, see e.g. Bostock *et al*., 2005; Bucher & Goaillard, 2011; Debanne *et al*., 2011; Kullmann *et al*. 2005). Short distances between nodes of Ranvier in group Ia afferents, about 300 µm in the cat (Ishizuka *et al*., 1979) and <200 and even <100 µm in group Ia and skin afferents in the rat (Lucas-Osma *et al*., 2018) within the entry zone of the afferents to the spinal grey matter may make the propagation of action potentials particularly vulnerable to branch point failures. When the activation of single fibres was analysed in an in vitro spinal cord preparation they were found to occur in around 20% of fibres (see Fig 2f of Hari *et al*., 2022). Indications for frequent branch point failures were also reported for monosynaptic EPSPs evoked in mice motoneurons and EMG responses in freely moving mice (see Fig.5 and 6 in Hari *et al*., 2022) and the monkey (Mahrous et al. 2024). The delays in propagation of action potentials appeared to be of the order of 0.1-0.2 ms over one internodal distance (see extended Figs. 2, 3, 5 & 6 in Hari *et al*., 2022) and to be reduced or eliminated by both GABA and fibre depolarization. The shortening in the delays of nerve volleys evoked by epidural stimulation during the post-polarization period (estimated to be 0.03-0.15 ms, our Fig 5) is well in keeping with these data. Epidural polarization may thus synchronize and speed up the monosynaptic actions of afferent fibres on their target cells and thereby render them more efficient, whether evoked by electrical or natural stimuli. As discussed previously (Jankowska & Hammar, 2021), reducing the conduction time along fibres providing input to spinal motoneurons and interneurons by 0.1 -0.2 ms may appear trivial and of negligible consequence. However, as nerve impulses reach their target neurons via several collaterals of the same as well as of different sensory fibres, and at slightly different latencies, a synchronization of their arrival together with the ephaptic coupling between afferent fibres prior to their entry into the spinal cord (Bolzoni & Jankowska, 2019) may increase the probability of activating their target neurons. Furthermore, a difference of 0.1-0.2 ms in conduction time along the presynaptic nerve fibres constitutes a non-negligible part of the rise time of about 1 ms of composite monosynaptic EPSPs in motoneurons and especially of unitary EPSPs evoked by single Ia afferents (0.25-0.60 ms in the fastest unitary EPSPs; Jack *et al*., 1971). The elimination of a sizeable proportion of branch point failures together with the increased conduction velocity in fibres entering the spinal grey matter would thus have a significant impact on the activity of spinal neuronal networks.

### The hypothesized spinal network modulating input to the spinal cord at the level of entry of the sensory information to the grey matter

Each of the components of the simplified diagram of the proposed network depicted in Fig. 2 is based on experimental data but we need to know much more about both the GABAergic interneurons and glial cells affecting the branching regions of the afferents to evaluate their relative importance. The probability that GABAergic interneurons in the deeper part of the dorsal horn contribute to this modulation has recently increased significantly. Particularly strong candidates for such neurones are GABA_axo_ neurons in GAD2//tdTOM//CHR2-EYFP mice (Hari *et al*., 2022, see also Gradwell, 2024) and some V3 neurones that provide input to them, as they were found to mediate the optogenetically evoked depolarization of afferent fibres (Hari *et al*., 2022; Lin *et al*., 2023) However whether the same or different GABAergic interneurons contribute to effects evoked via axo-axonic contacts or via volume transmission of GABA. has as yet not been resolved.

The GABAergic interneurons in the intermediate zone would likely be activated by group I afferents and mediate axo-axonally induced depolarization of group I afferents. Interneurons in laminae III-IV would, on the other hand, more likely be activated by group II and low threshold cutaneous afferents, especially as interneurons at such location were found to induce spike-triggered DRPs in feline sacral segments (Jankowska & Riddell, 1995) and include not only glutamatergic and glycinergic but also GABAergic interneurons (Gradwell, 2024) If interneurons with input from group II and skin afferents constitute the main source of GABA at the level where afferent fibres enter the grey matter, the tonic input from muscle spindle secondaries and skin would allow them to provide a background level of GABA close to the proximal branching regions of the afferents. Under conditions of the present experiments such tonic input would be provided by proximal branches of the sciatic nerve and by the femoral nerve but in intact animals by any nerves.

We cannot estimate the extent to which the effects of DC delivered at the site of afferent entry to the spinal cord reflect unspecific effects of the extrasynaptic GABA spill-over (by either glial cells or GABAergic interneurons) and how these are related to selective actions of GABAergic interneurons evoked via axo-axonic synaptic contacts within more distal compartments of the fibres. Hari *et al*. (2022) and Bradesi *et al*. (2010) considered the possibility of specific actions of GABA both when it acts via axo-axonic contacts and via volume transmission, concluding that “…GABA near sodium channels provides a powerful mechanism to turn on specific nodes and branches to regulate sensory feedback.” (Lin *et al*., 2023 p.1289 left column). However, it has not yet been established whether only GABAergic interneurons or both these interneurons and the glial cells are the sources of the optogenetically released GABA and whether GABA from these two sources acts within the same branching regions. If not, GABA originating from glial cells and acting on branching sites closest to or within the dorsal columns might have generalized and unspecific facilitatory effects while GABAergic interneurons affecting more distal branching sites on axon collaterals close to their target cells might be more selective and specific. The generalised effects of extrasynaptically acting GABA at the level of entry of the spinal afferent fibres might be reminiscent of …” setting the overall excitability of the system….. by a homeostatic regulation of the overall level of GABA released into the extracellular space ”…. Kullmann *et al*., 2005 p.8) in the central nervous system.

Assuming that glial cells within the most dorsal regions of branching of afferent fibres increase the extracellular concentration of GABA both when depolarised by epidurally applied DC and by afferent fibres (Christensen *et al*., 2018), it would be also important to know whether afferent input to glial cells and to GABAergic interneurons releasing GABA within the extracellular space differs from input to interneurons forming axo-axonic contacts. It would also be of great interest to compare afferent input to excitatory interneurons that relay it to GABAergic interneurons (Lin *et al*., 2023) and to glial cells (Christensen *et al*., 2018) and to know whether both AMPA and NMDA membrane receptors and both VGlut1 and VGlut2 contacts are present on proximal as well as distal nodes, .It might be expected that actions of GABA released into the extracellular space at the interface between the dorsal columns and the dorsal horn, where axon collaterals of all categories of afferent fibres are mixed, are general rather than specific. The proximal effects of GABA released by glial cells might accordingly facilitate synaptic actions of muscle afferents on motoneurones in several motor nuclei and affect both muscle and skin afferents targeting various populations of spinal interneurons. The effects of DC on the excitability of different categories of afferent fibres have as yet not been systematically compared. Nevertheless, it has been a common finding that both early and later components of nerve volleys evoked by epidural stimulation and attributed to group I and group II muscle afferents are affected by DC. In the presently reported results, this is illustrated in Fig. 1 C and Fig. 5A and C showing that both components of these volleys are not only increased but also shifted to the left. The DC-evoked increase in synaptic actions of afferent fibres was likewise seen not only in motor nuclei (Bączyk *et al*., 2019; 2020) but also in the dorsal horn (field potentials from group II and /or skin afferents (Bączyk & Jankowska, 2018).

Despite all these unanswered questions the networks hypothesized in Fig. 2 appear to represent the basic system gating the level of input to the spinal cord partly by the glial cells and partly by sensory feed-back via GABAergic interneurons.

## Authors contribution

Both authors contributed to the study design, data collection and interpretation, and drafting the manuscript. Data analysis and preparation of the figures by EJ.

## Acknowledgments

We wish to thank S. Berg and D. Magnusson for excellent technical support an our colleagues for discussions and helpful comments on earlier versions of the manuscript. The study was supported by the Institute of Neuroscience and Physiology.

## Data accessibility

The reported experimental data are available on request from the corresponding author.

## Abbreviations

DC: direct current
ECG: electrocardiogram
EMG: electromyography
GABA: γ-Aminobutyric acid
L: Lumbar
L-AAA: L-alpha-aminoadipic acid
PAD: primary afferent depolarisation
Per: peroneal nerve
T: threshold
Tib: tibial nerve

